## Supplementary material for "Characterization and clustering of kinase isoform expression in metastatic melanoma": Table S2

***Expanded RTK table***

**Table S2:** RTKs with significant differential expression between high purity primary tumor and metastatic samples.

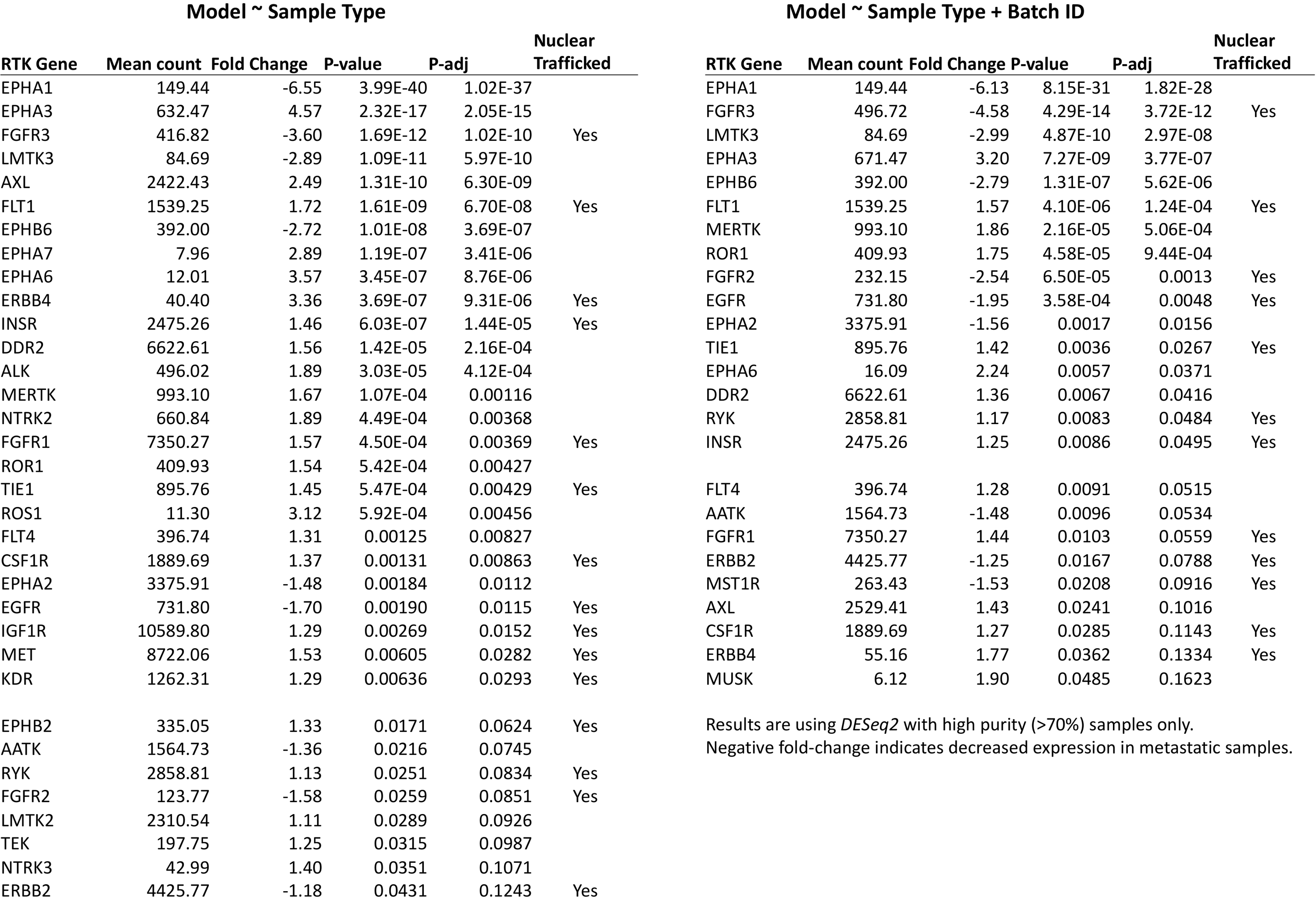
