## Supplementary material for "Characterization and clustering of kinase isoform expression in metastatic melanoma": Figure S1

***Effects of sample impurity and 3’ fragment bias on apparent DIR***

To test the effects of data bias on apparent DIR significance, we removed samples one-by-one in order of either highest impurity or highest 3’ bias. P-values were calculated (PCA test) at each iteration. Samples were then removed in reverse order for comparison. Shown in Figure S1 are the results for one affected gene, *EIF2AK4*. Significance was driven by high levels of the 3’ fragment *EIF2AK4-205* (Figure S1A) in some primary tumor samples (Figure S1B,C). These artefacts biased the interpretation of significant differences in isoform expression (Figure S1D,E) and fragment bias accounted for the bulk of the low p-values observed. For a few genes, sample impurity was the culprit. For example, levels of *PTK2B* (isoform-205) were much higher in immune infiltrate samples (Figure S2).

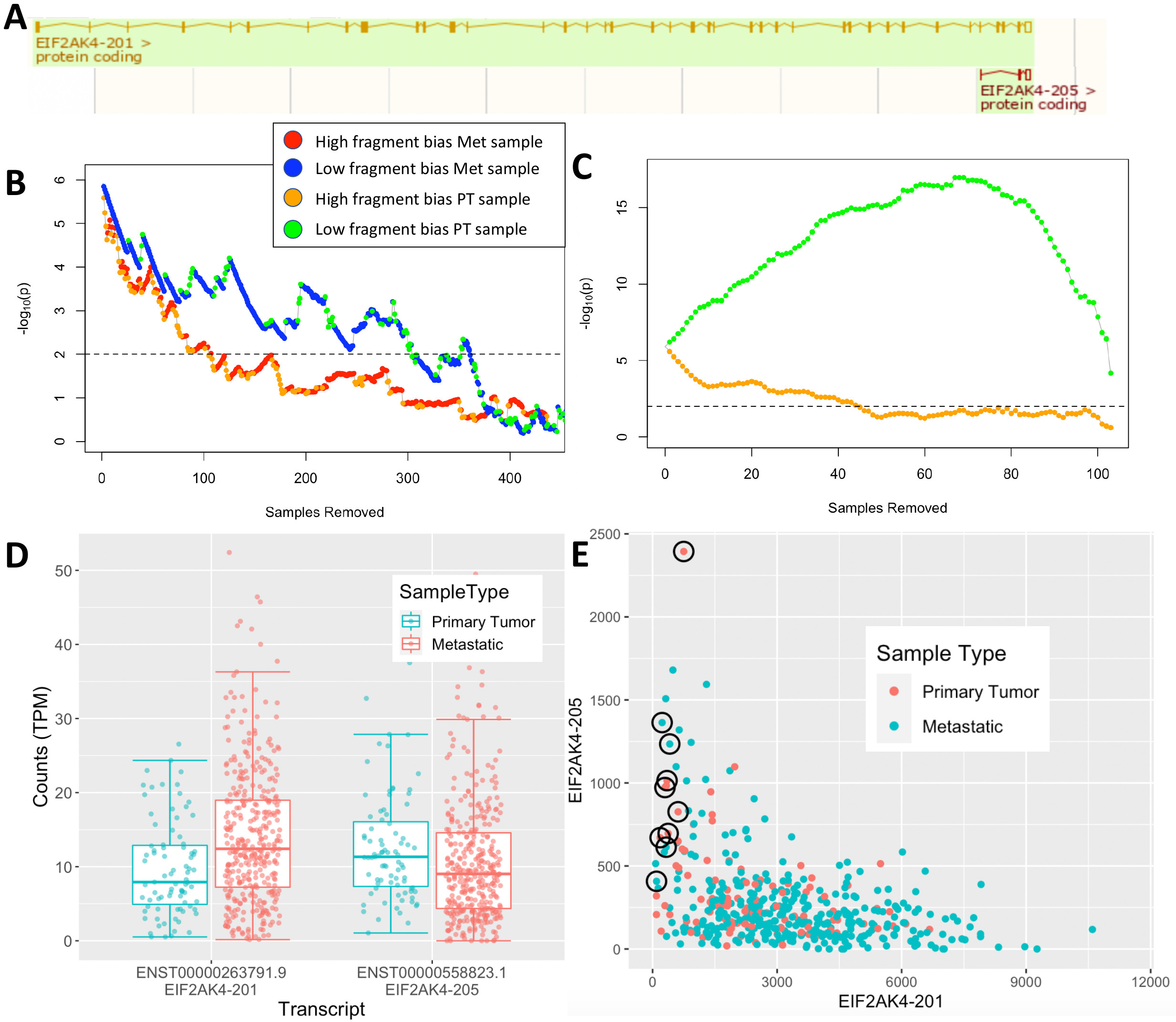

**Figure S1: Fragment bias in primary tumor samples drives significance in EIF2AK4.** **(A)** Two protein coding isoforms of EIF2AK4, the full-length isoform (-201) and 3’ fragment (-205). **(B)** Change in DIR significance (-log_10_p) as samples are removed one-by-one in order of highest bias (red and orange dots) vs in order of lowest bias (green or blue dots). The significance drops faster when the high-bias samples are removed. The p-value here is calculated using the PCA method with the coin general independence test. (C) When only primary tumor samples are removed, the difference in p-values are even more disparate, indicating that high-bias primary tumor samples drive significance. (D) Box plots for the three isoforms with the highest number of normalized counts. Significance is driven by a higher amount of the full-length isoform in metastatic samples but a lower amount (on average) of the 3’ fragment. (E) Scatter plot of the raw counts of each isoform in each sample. Circled in black are the ten isoforms with the highest 3’ bias, indicated by high levels of the 3’ fragment and low levels of the full-length isoform. I would remove EIF2AK4-210 since it is not shown in 2A.
