## Supplementary material for "Characterization and clustering of kinase isoform expression in metastatic melanoma": Figure S2

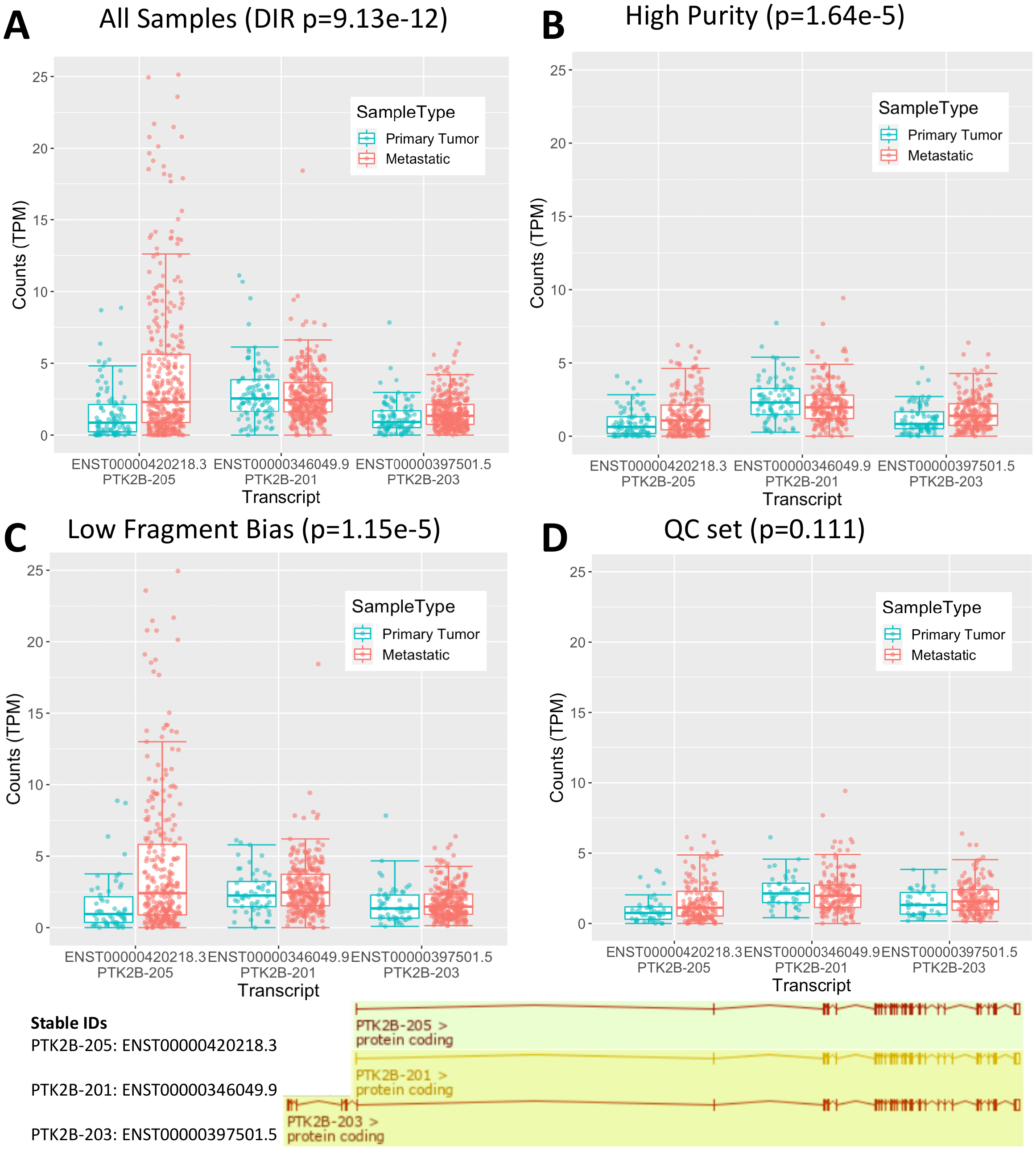

**Figure S2. Quality control reduces DIR significance of PTK2B.** Expression of isoform PTK2B-205 in particular is driven by low-purity metastatic samples. Its expression drastically decreases when they are removed. Conversely, there is lower average expression of PTK2B-203 in primary tumor samples before samples with high 3’ bias are removed. This is likely due to the presence of more exons on the 5’ end, which will be undercounted in samples with 3’ bias.
