## Supplementary material for "Characterization and clustering of kinase isoform expression in metastatic melanoma": Figure S3

***Genes with both DE and DIR***

Some genes have DIR which coincides with significant DE. For example, 6 of the 7 coding isoforms of *FGFR3* are suppressed in metastatic samples (Figure S3), while the remaining isoform -205 has mildly increased expression, altering isoform ratios. *PAK6*, with 14 isoforms, undergoes a similar alteration. In *BLK*, DIR of 3 isoforms is driven by an unequal increase of 2 major isoforms, rather than all 3.

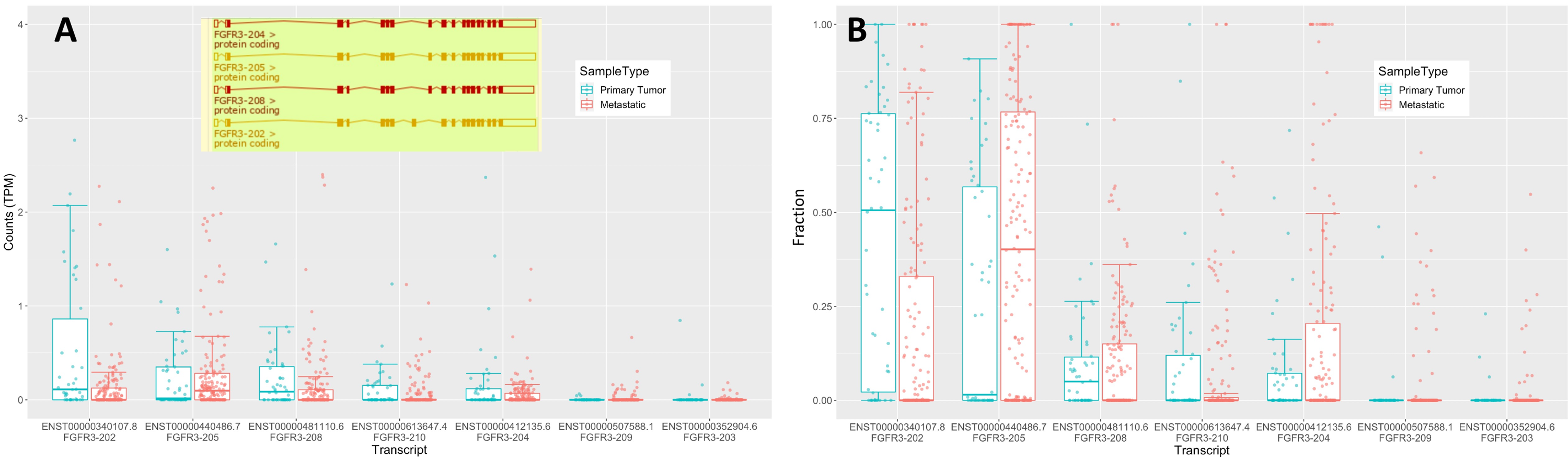

**Figure S3. Differential isoform ratios in FGFR3**. Plotted are (A) TPM counts and (B) fraction of all isoform counts for each sample. Although the trend is decreased expression, one isoform (FGFR3-205) has mildly increased expression, resulting in highly altered isoform ratios.
