## Supplementary material for "Characterization and clustering of kinase isoform expression in metastatic melanoma": Figure S4

***Isoform expression clusters***

The k-means elbow method identified 4 isoform groups and 4 sample clusters (Figure S4). Group 1, the largest group, was enriched in isoforms for genes involved in “blood vessel development” (one-sided Fisher’s exact test, p=7.4e-5), “MAPK cascades”, and “positive regulation of cell differentiation”. Group 2, which was highly expressed in low-purity samples, was enriched for genes in involved in immune response. Group 3, highly expressed in Cluster C (which is enriched for *RAS* mutants) contains isoforms for genes that regulate cell motility. The remaining group, which contains several isoforms with correlated downregulation, was enriched for “positive regulation of transcription”

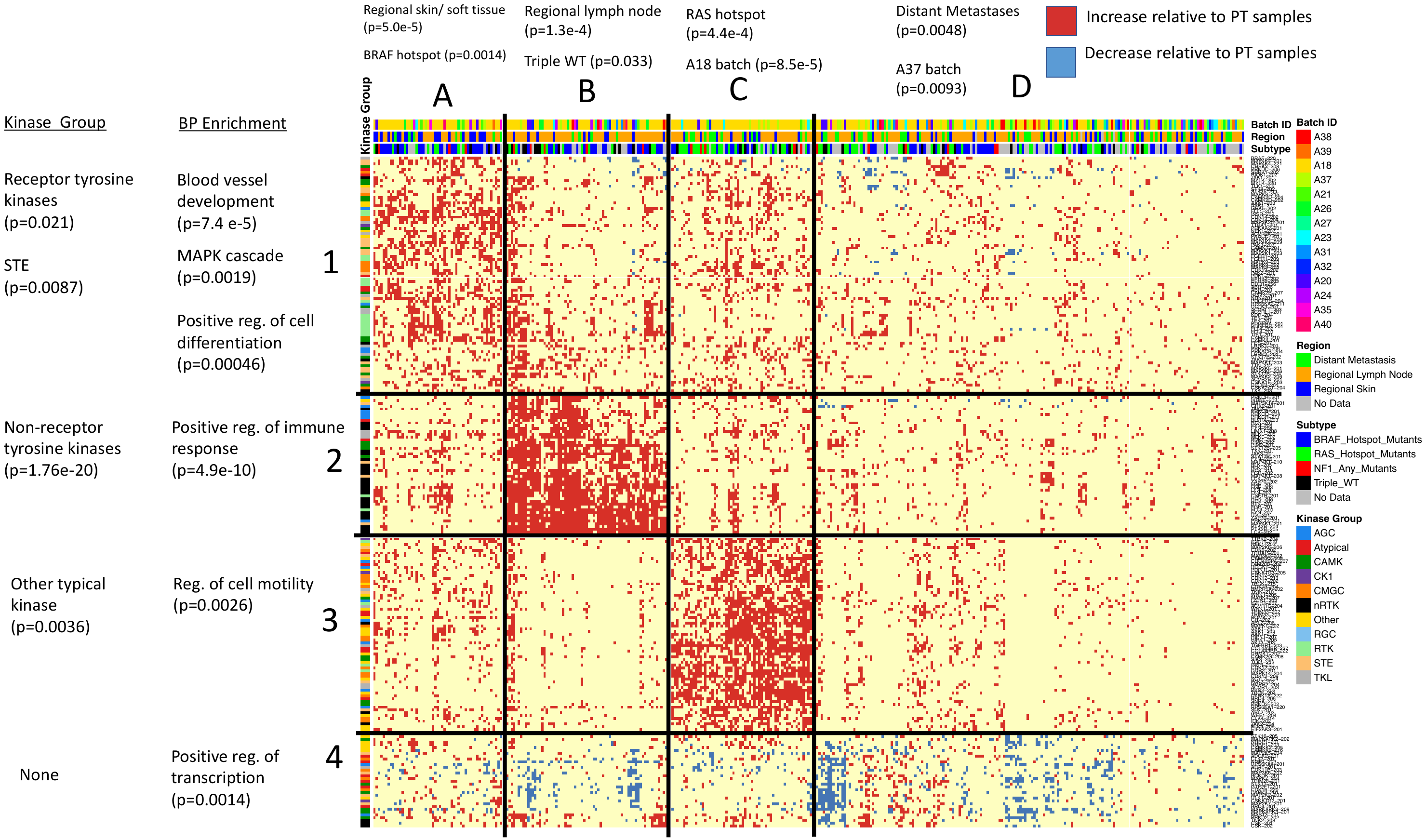

**Figure S4:** **Heatmap of 367 metastatic samples clustered (4x4) according to kinase isoform counts.** Red dots indicate increased expression in metastases (Quasi-Poisson GLM, p<0.05) while blue dots indicate decreased expression (p<0.2). Shown are the 367 metastatic samples (columns) and 235 isoforms that were altered in >13% of samples (rows). P-values were calculated using Fisher’s exact test.

However, we found that clustering using 5 isoform groups enhanced certain enrichment patterns, and we discuss the 5x4 results in the main paper.
