## Supplementary material for "Characterization and clustering of kinase isoform expression in metastatic melanoma": Figure S5

Because Cluster C was also enriched for batch A18, we separated the samples into two groups: batch A18 samples (n=199) and all other samples (n=168), and re-clustered the samples in both subsets (Figure S5). We found that both subsets of samples separate into four clusters comparable to those shown in Figures 8 and S3. Furthermore, the 3^rd^ cluster is still enriched for *RAS* mutants into both subsets; it is just that the A18 batch contains more *RAS* mutants relative to the other batches. Thus the enrichment pattern of Cluster C is independent of batch.

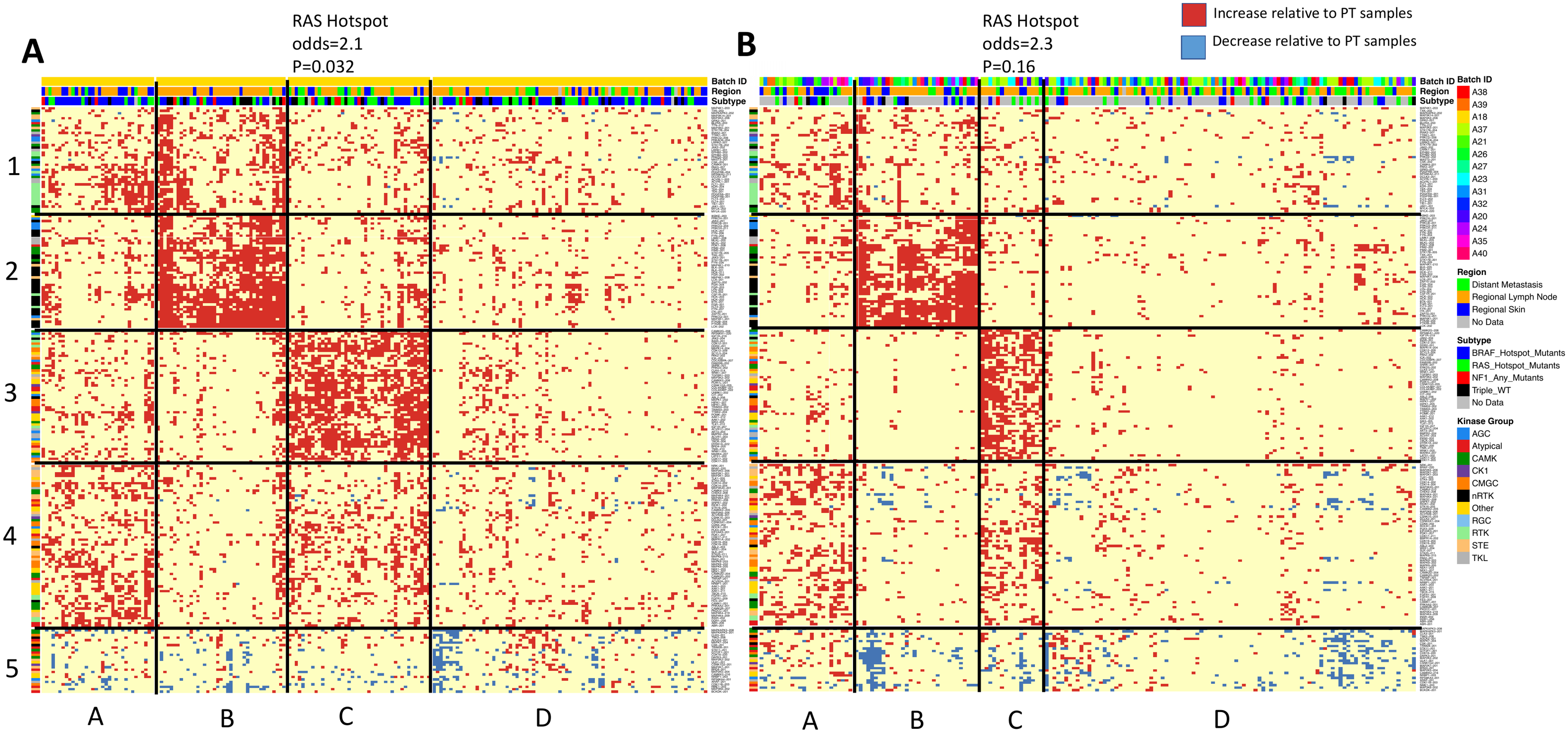

**Figure S5:** **Clustering of batch A18 and non-batch A18 samples.** Isoform groups are the same as in Figure 8. Both sample subsets separated into four clusters comparable to Figures 8 and S3.
