## Supplementary material for "Characterization and clustering of kinase isoform expression in metastatic melanoma": Figure S6

***Apoptosis in A375 melanoma cells with heightened* SLK *isoform expression***

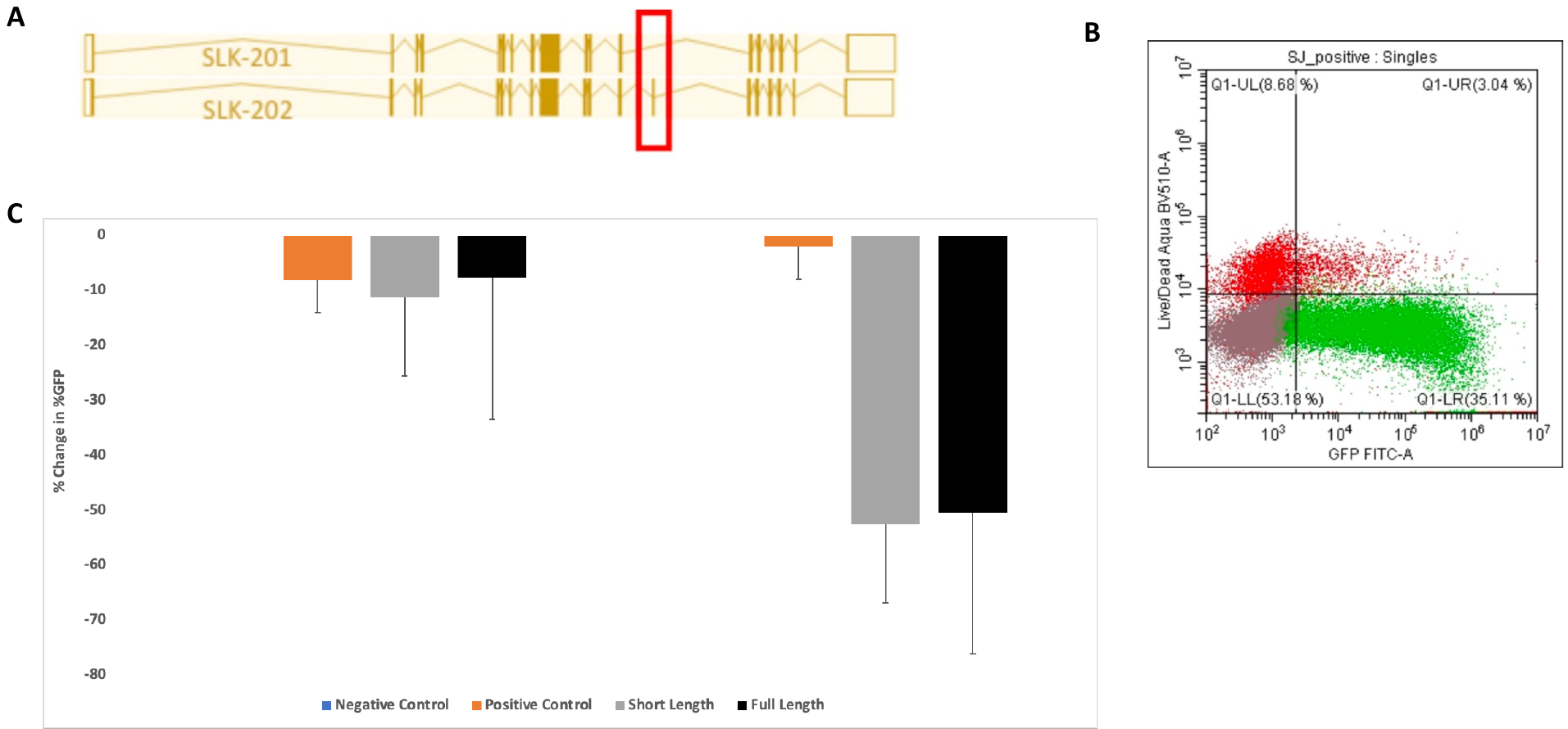

**Figure S6. SLK isoform expression results in cell death (A)** This schematic shows the structure of the two different SLK isoforms. Short-length SLK, SLK-201, codes for a shorter isoform due to a missing exon. Full-length SLK, SLK-202, is not missing any exons. We hypothesize that transient overexpression of SLK full-length-GFP will produce more cell death than SLK short-length-GFP in metastatic melanoma.  **(B)** Here we show a representative FACs output thresholded by live/dead cells and GFP. We analyzed the % change in % GFP over 72h for the negative control, positive control (empty eGFP fusion vector), short-length SLK-GFP, and full-length SLK-GFP. **(C)** A bar graph showing the results of our time course analysis. We see a significant reduction in % change of %GFP for both SLK isoforms between 48h and 72h in live cells that is not seen in the positive control. This suggests that both SLK isoforms cause cell death between 48h and 72h. No significant difference in % change in %GFP was observed between the short length and the full-length isoform indicating that they produce about the same amount of cell death.
