## Supplementary material for "Characterization and clustering of kinase isoform expression in metastatic melanoma": Figure S7

***Heightened expression of* BRD4 *isoforms in* RAS*-mutant metastatic samples***

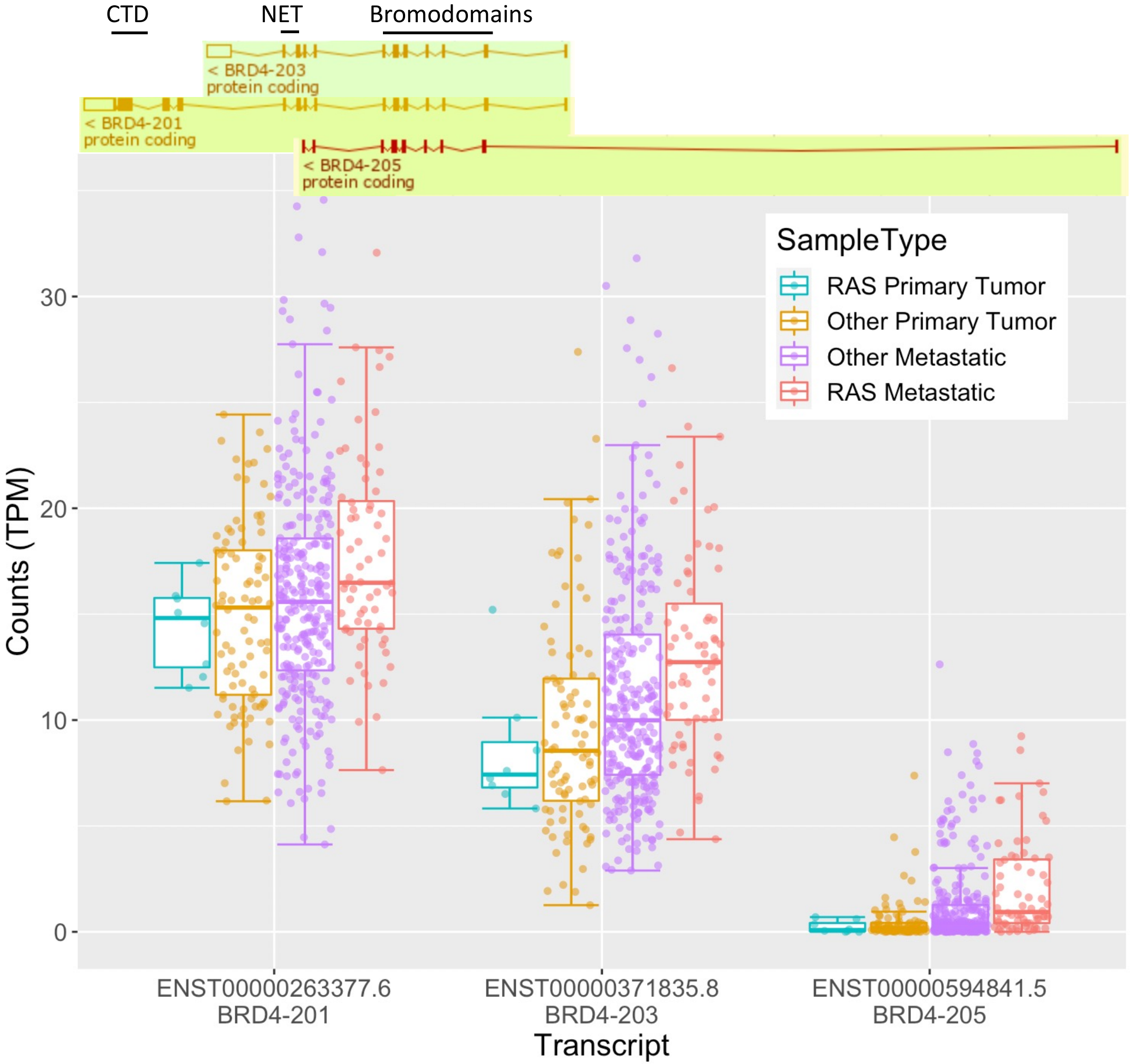

**Figure S7. Heightened expression of BRD4 isoforms in RAS mutant metastatic samples.** Although total BRD4 counts did not test as having significant DE between any group of primary and metastatic tumors, isoforms BRD4-203 and BRD4-205 have heightened expression in RAS-mutant metastatic samples. Exon junction analysis did not confirm these particular isoforms.
