## Supplementary material for "Characterization and clustering of kinase isoform expression in metastatic melanoma": Empirical FDR

***Empirical FDR adjustment of biological process (BP) annotation enrichment***

Although we choose not to adjust p-values for BP enrichment (see Methods in main paper), we did calculate an adjusted p-value “q” based on an empirical false discovery rate:

$$E[FDR(p)]=\frac{E[V\left( p \right)]}{R(p)}$$

Where E[V(p)] is the expected number of significant annotations (discoveries) from a random set of kinase genes at threshold “p”, averaged from 1,000 iterations, and R(p) is the total number of discoveries at threshold “p” from a set of results. For ranked p-values, R(p_i_) = i for a p-value at rank “i”. As with Benjamini-Hochberg adjustment, the q-value at rank “i” is the minimum expected FDR for all annotations greater than or equal to rank “i”.

$$q_{p_{i}}=\min_{j\in[i,n]} \{E\left[ FDR\left( p_{j} \right) \right] \}$$

Where “n” is the number of tested annotations and thus the max rank.

A high “q” does not mean a set of results should be discarded. On the contrary, lack of enrichments indicates significant genes are dispersed across many biological processes rather than concentrated. Below are the lowest q-values obtained for each set of results from Tables 3 and 5 in the main paper. Although q-values >1 are typically set to 1 (a true FDR cannot exceed 1 in a single experiment), here we chose to keep q as calculated to indicate lack of enrichments.

**Minimum Q-values of BP enrichments for analyzed sample sets**

| **Analysis** | **Sample Set** | **Strongest BP enrichment for top 5% of genes** | **P-value** | **Q-value** |
| --- | --- | --- | --- | --- |
| DE | All samples | Adaptive immune response | 5.09e-9 | **<6.67e-5** |
|  | HP samples | Regulation of lymphocyte activation | 0.024 | 7.70 |
|  | BRAF mutant HP | Cell differentiation | 1.3e-4 | **0.036** |
|  | RAS mutant HP | Eye morphogenesis | 0.0034 | 2.08 |
|  | NF1 mutant HP | Regulation of MAPK cascade | 0.0054 | 0.64 |
|  | Triple wildtype HP | Intracellular receptor  signaling pathway | 9.7e-4 | 0.65 |
| DIR | All samples | Positive regulation of translation | 1.5e-4 | **0.043** |
|  | QC samples | Regulation of endocytosis | 3.5e-4 | 0.23 |
|  | BRAF mutant QC | Regulation of protein acetylation | 0.0084 | 1.26 |
|  | RAS mutant QC | Positive regulation of angiogenesis | 1.6e-4 | **0.048** |
|  | Triple wildtype QC | protein modification by small protein conjugation or removal | 0.0027 | 1.46 |
